# Susceptibility to Zika virus in a Collaborative Cross mouse strain is induced by *Irf3* deficiency *in vitro* but requires other variants *in vivo*, not linked to the type I interferon response

**DOI:** 10.1101/2023.03.27.534491

**Authors:** Marie Bourdon, Caroline Manet, Laurine Conquet, Corentin Ramaugé-Parra, Etienne Kornobis, Eliette Bonnefoy, Xavier Montagutelli

**Affiliations:** Institut Pasteur, Université de Paris, Mouse Genetics Laboratory, F-75015 Paris, France; Institut Pasteur, Université de Paris, Biomics, F-75015 Paris, France; Université Paris Cité, Institut Cochin, Inserm, CNRS, F-75014 Paris, France

**Keywords:** Zika virus, mouse, Irf3, interferon response, Collaborative Cross

## Abstract

Zika virus (ZIKV) is a Flavivirus responsible for recent epidemics in Pacific Islands and in the Americas. In humans, the consequences of ZIKV infection range from asymptomatic infection to severe neurological disease such as Guillain-Barré syndrome or fetal neurodevelopmental defects, suggesting, among other factors, the influence of host genetic variants. We previously reported similar diverse outcomes of ZIKV infection in mice of the Collaborative Cross (CC), a collection of inbred strains with large genetic diversity. CC071/TauUnc (CC071) was the most susceptible CC strain with severe symptoms and lethality. Notably, CC071 has been recently reported to be also susceptible to other flaviviruses including dengue virus, Powassan virus, West Nile virus, and to Rift Valley fever virus. To identify the genetic origin of this broad susceptibility, we investigated ZIKV replication in mouse embryonic fibroblasts (MEFs) from CC071 and two resistant strains. CC071 showed uncontrolled ZIKV replication associated with delayed induction of type-I interferons (IFN-I). Genetic analysis identified a mutation in the *Irf3* gene specific to the CC071 strain which prevents the protein phosphorylation required to activate interferon beta transcription. We demonstrated that this mutation induces the same defective IFN-I response and uncontrolled viral replication in MEFs as an *Irf3* knock-out allele. By contrast, we also showed that *Irf3* deficiency did not induce the high plasma viral load and clinical severity observed in CC071 mice and that susceptibility alleles at other genes, not associated with the IFN-I response, are required. Our results provide new insight into the *in vitro* and *in vivo* roles of *Irf3*, and into the genetic complexity of host responses to flaviviruses.

**Author summary:** Recent ZIKV outbreaks led to millions of infected people, with rare but severe complications such as Guillain-Barré syndrome and encephalitis in adults suggesting that host genes influence the susceptibility to severe forms of infection. We previously reported the importance of host genes in ZIKV pathogenesis using a panel of genetically diverse mouse strains and identified CC071 as the most susceptible strain. Importantly, this mouse strain has been shown by others to be also susceptible to several other RNA viruses. Through a combination of functional and genetic approaches in a cellular model, we identified a mutation in the *Irf3* gene which plays a key role in activating the expression of interferon beta to induce the type I interferon response, the first line of host defense against the virus. This mutation fully explains the high viral replication observed in CC071 cells. However, it was not able to induce the elevated viremia and the symptoms displayed by CC071 ZIKV-infected mice, unraveling the implication of other host genes which are not associated with the type I interferon response. Because of the broad susceptibility of CC071 to multiple viruses, our results have implications beyond ZIKV infection and contribute to shedding light on the plurality of host mechanisms fighting infectious diseases.

## Introduction

Zika virus (ZIKV) is a mosquito-borne virus of the *Flaviviridae* family identified in 1947 in Uganda. The first noticeable human outbreaks occurred in Micronesia in 2007 and in French Polynesia and New Caledonia in 2013-2014. In 2015-2016, ZIKV caused an epidemic in Brazil which rapidly spread across the Americas and the Caribbean. To date, 89 countries have reported evidence of mosquito-transmitted Zika virus infection (https://www.who.int/health-topics/zika-virus-disease#tab=tab_1).

While most people infected with ZIKV remain asymptomatic, some develop non-specific symptoms including rash, fever, conjunctivitis, muscle and joint pain, malaise and headache. Neurological complications have been described in adults such as Guillain-Barré syndrome (1) and encephalitis (2). Infection of pregnant women was associated with congenital Zika syndrome in the fetus, which can lead to neurodevelopmental deficiencies, brain malformation (3) or in some cases to fetal loss (4).

Many factors may contribute to this variable severity, including the viral strain, the infection route and dose, and the host genetic background (5,6). Indeed, mouse and human studies have shown that host genes influence flaviviral infections’ outcomes (7). While human genetic studies are hampered by the variability of these multiple factors, they can be controlled in mouse models which have proven very valuable to identify susceptibility variants (8,9). Relevant ZIKV infection models have been developed in mice either using *Ifnar1* knock-out (KO) mice in which the IFNAR receptor to IFN-I has been inactivated (10,11), or by blocking this receptor using a monoclonal antibody targeting the IFNAR1 receptor subunit (MAR1-5A3 (12)).

We have previously explored the role of mouse natural genetic variants on ZIKV susceptibility in the Collaborative Cross (CC), a panel of recombinant inbred mice encompassing a genetic diversity similar to that of the human population and capturing approximately 90% of the mouse natural genetic variants (13,14). We reported that the CC genetic diversity enabled large variations in the clinical severity of ZIKV disease, plasma viral load and intensity of brain pathology, comparable to those observed in human cases (15). We specifically identified CC071/TauUnc (CC071) mice as very susceptible, with high mortality and high peak plasma viral load. Notably, the CC071 strain has been recently reported as also susceptible to other flaviviruses including dengue virus (15), Powassan virus (16), West Nile virus (WNV) (15), and to Rift Valley fever virus (17), emphasizing its value to decipher genetic factors controlling host responses to RNA viruses.

We previously demonstrated that genetic background influenced ZIKV replication in CC mouse embryonic fibroblasts (MEFs). Here, we investigated the mechanisms driving high viral replication in CC071 MEFs. We found that, compared with the more resistant C57BL/6J (B6) and CC001/Unc (CC001) strains, ZIKV-infected CC071 MEFs displayed a delayed expression of the IFN-I response genes. Genetic and functional analyses identified a strain-specific variant in the *Irf3* gene, the first transcription factor involved in interferon (IFN) β expression, which mimics the effects of a null allele *in vitro* and fully explains the delayed IFN-I expression and uncontrolled viral replication. By contrast, we showed, from the *in vivo* comparison of CC071, *Irf3*-deficient and backcross mice, that the CC071 *Irf3* mutation is not sufficient to explain the high susceptibility of CC071 mice to ZIKV infection and that other genes, not associated with the IFN-I response, are involved. These findings provide new insights into the roles of *Irf3* in viral diseases and exemplify how the study of CC strains allows deciphering the role of host genes in viral pathogenesis.

## Results

### CC071 MEFs show defective control of viral replication and delayed IFN-I expression, but normal response to IFN-I stimulation

We previously reported that, unlike CC001 MEFs, CC071 MEFs produced increasing quantities of viral particles during the first 72 hours post-infection (hpi) (15). Here, we confirmed and expanded this observation by infecting B6, CC001 and CC071 MEFs and by quantifying viral particles by FFA. After ZIKV infection, CC071 MEFs displayed high and increasing viral titers between 24 and 72 hpi, while CC001 and B6 MEFs showed stable and lower titers (Fig 1A). To investigate the origin of the defective control of viral replication in CC071 MEFs, we measured the expression level of the *Ifnb1* gene coding for IFNβ in ZIKV-infected CC071, CC001 and B6 MEFs. *Ifnb1* expression is induced very rapidly after virus detection by sensors and triggers the innate antiviral response which is essential for limiting viral replication. In CC001 and B6 MEFs, *Ifnb1* expression was significantly induced at 24 hpi and remained stable and high until at least 72 hpi (Fig 1B). In contrast, its expression in CC071 MEFs was low at 24 hpi and reached the level of CC001 only at 72 hpi. Similar results were obtained for *Ifn4a* which encodes one of the IFNα proteins (Sup Fig 1). Notably, *Ifnb1* expression in CC071 MEFs at 72 hpi was significantly higher than in B6 MEFs, showing that CC071 MEFs were delayed but not intrinsically hampered in their ability to induce strong IFN-I expression.

**Figure 1.**
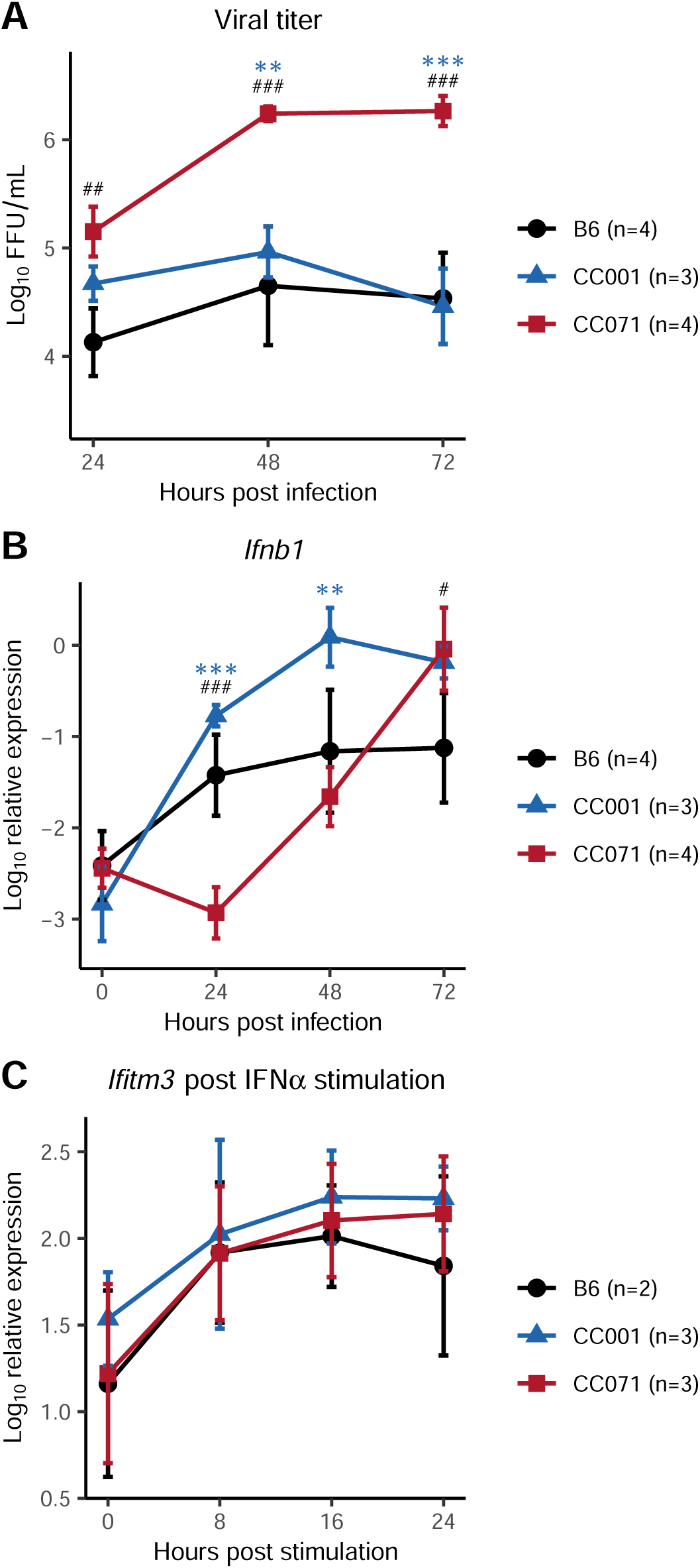
CC071 MEFs fail to control viral replication, with delayed *Ifnb1* expression but normal response to IFN-I. (A, B) MEFs derived from B6 (black circles), CC001 (blue triangles) and CC071 (red squares) were infected with ZIKV at a MOI of 5 and analyzed 24, 48 and 72 hpi. (A) Viral titer in supernatants was quantified by FFA. (B) *Ifnb1* expression was normalized to *Tbp* reference gene. Data are mean +/- sem from 3 to 4 biological replicates per strain (MEFs derived from individual embryos). (C) MEFs were stimulated with recombinant IFNα. *Ifitm3* relative expression normalized to *Tbp* reference gene is shown as an example of ISG. Data are mean +/- sem from 2 to 3 biological replicates. Blue asterisks and black hashes show statistical significance of CC071 compared to CC001 and to B6, respectively (ANOVA followed by Tukey HSD, */# p < 0.05, **/## p < 0.01, ***/### p < 0.001).

To test whether this defective induction of *Ifnb1* expression was specific to ZIKV infection, MEFs were then transfected with the influenza A virus-derived 3-phosphate-hairpin-RNA (3p-hpRNA), an agonist of the RIG-I/MDA5-MAVS pathway, or treated with polyinosine-polycytidylic acid (poly (I:C)), that activates both Toll-like receptor (TLR) 3 (TLR3) and the RIG-I/MDA5-MAVS pathway (18). Here again, CC071 MEFs showed a delayed expression of *Ifnb1* after both stimulations by comparison with B6 and CC001 MEFs (Sup Fig 2), indicating that the defect in IFN-I genes expression in CC071 MEFs was not specific to ZIKV infection. This result suggested a defect in the molecular cascade between cellular sensors of pathogen-associated molecular patterns (PAMP) and the *Ifnb1* gene transcription machinery.

**Figure 2.**
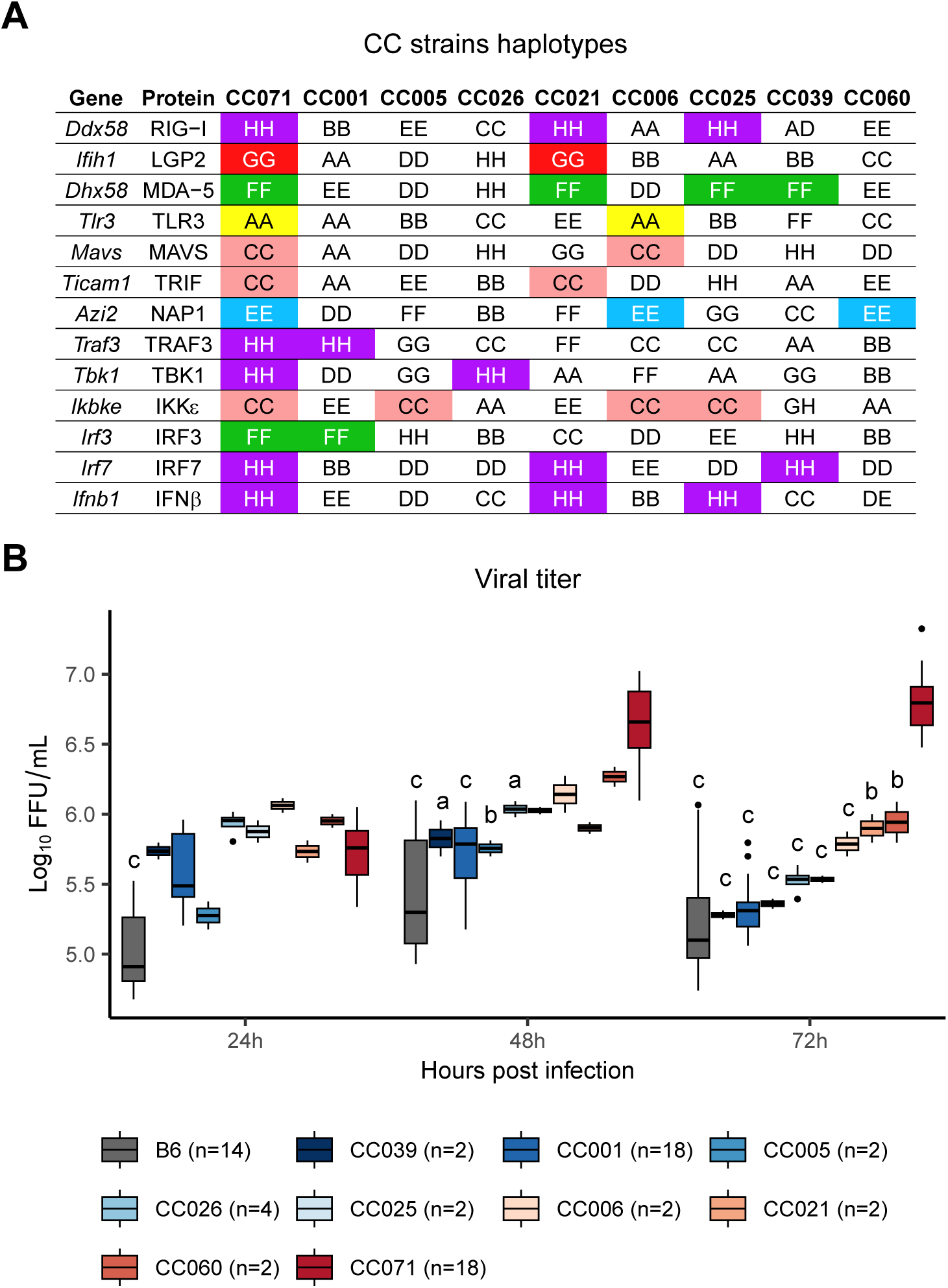
Haplotype analysis fails to identify a gene from the *Ifnb1* induction pathway associated with uncontrolled viral replication. (A) Identification of CC strains which carry the same ancestral haplotype as CC071 at the genes involved in the pathway leading to *Ifnb1* expression. Colored boxes indicate matched haplotypes between CC071 and other CC strains. Letters and colors designate the eight CC founder strains. A: A/J (yellow); B: C57BL/6J; C: 129S1/SvImJ (pink); D: NOD/ShiLtJ; E: NZO/HILtJ (light blue); F: CAST/EiJ (green); G: PWK/PhJ (red); H: WSB/EiJ (purple). Doubled letters (eg AA) indicate homozygous genotypes. Heterozygous genotypes are indicated by the two corresponding letters (eg AD). (B) Kinetics of viral titer in MEFs from CC071 and the 8 CC strains shown in (A). Experimental conditions were as in Figure 1A. n: number of technical replicates for each strain. Data are mean +/- sem from the technical replicates. Letters show statistical significance between CC071 and other strains. (ANOVA followed by Tukey HSD, a: p < 0.05, b: p < 0.01, c: p < 0.001).

To evaluate the capacity of CC071 MEFs to respond to IFN-I, they were treated with recombinant IFNα. The expression of IFN-stimulated genes (ISGs) such as *Ifitm3* was induced with the same kinetics and level as in B6 and CC001 MEFs (Fig 1C), showing that CC071 MEFs are able to respond normally to IFN-I stimulation and that their defect is limited to the induction of *Ifnb1* gene expression.

### CC071’s delayed Ifnb1 expression is strain-specific

To gain insight into the mechanisms responsible for defective *Ifnb1* induction, we investigated the expression levels of all genes involved in *Ifnb1* expression on CC071, B6 and CC001 MEFs at 16, 24 and 32 hpi. Mock-infected MEFs were analyzed at 24 hours as controls. Expression levels were measured by RNA sequencing (RNAseq) which provided a comprehensive analysis of transcriptomic changes. In CC001 MEFs, the expression of many genes rapidly increased after infection (160 at 16h, 821 at 24h and 971 at 32h; log2 fold-change > 1, FDR = 0.05), reflecting a robust innate antiviral response (Sup Fig 3A). A similar pattern was observed in B6 MEFs. By contrast, the expression of only 38 genes was increased in CC071 MEFs at 32hpi (34 of which were also activated in CC001), consistent with the delayed induction of *Ifnb1* expression. Among the genes that are involved in the pathway between PAMP sensors and *Ifnb1* transcription, ISGs such as *Tlr3*, *Ddx58* (coding for RIG-I sensor) or *Irf7* were not activated upon infection in CC071, while constitutively expressed genes such as *Mavs*, *Ticam1* (coding for the TRIF adaptor), *Traf3* or *Irf3*, showed comparable levels of expression in the three strains (Sup Fig 3B). Therefore, this analysis did not provide new clues for identifying the gene responsible for the defect observed in CC071.

We then leveraged the genetic architecture of the CC which genomes are patchworks of haplotypes inherited from the eight founder strains (19). Although CC071 was the only strain with severe ZIKV disease, we hypothesized that, if the delayed activation of *Ifnb1* resulting in uncontrolled viral replication observed in CC071 MEFs was due to an allele at one of the genes involved in the *Ifnb1* induction pathway inherited from a parental strain, ZIKV-infected MEFs of CC strains carrying the same allele would present similarly high viral titers. We therefore derived MEFs from each CC strain available to us carrying the same ancestral haplotype as CC071 at one of the 13 genes of the pathway (Fig 2A). Upon ZIKV infection, none of these CC MEFs showed viral titer kinetics resembling that observed in CC071 MEFs (Fig 2B). These results suggested two alternative hypotheses. Either the delayed *Ifnb1* activation involved two members of the pathway with a CC071-specific allelic combination leading to a non-functional interaction, or CC071 was carrying a strain-specific allele at one of these genes, resulting from a mutation proper to CC071 that probably arose on an ancestral haplotype during the CC071 inbreeding. However, the sequencing of one male of each CC strain reported in 2017 (20) did not identify such “private” variants with high predicted impact in CC071 for any of these genes. Whatever the molecular mechanism, our results indicated that it was specific to CC071.

### Genetic analysis identifies Irf3 as a candidate gene in a haplotype shared between CC071 and CC001

We then turned to a genetic mapping approach. We first established that (CC001xCC071)F1 MEFs responded to infection with as rapid induction of *Ifnb1* expression as CC001 MEFs (Sup Fig 4), suggesting that this CC071 trait was recessively inherited. F1 mice were therefore backcrossed with CC071. MEFs were produced from each of 51 backcross (BC) embryos, infected with ZIKV, and analyzed for viral titer and *Ifnb1* expression as above. One CC001 and one CC071 MEF lines were included in each infection experiment as controls. BC MEFs displayed either rapid and high *Ifnb1* expression with low viral titer (like CC001), or high viral titer and delayed *Ifnb1* expression (like CC071, Fig 3A-B), showing that these two traits correlated across the BC diverse genetic backgrounds.

**Figure 3.**
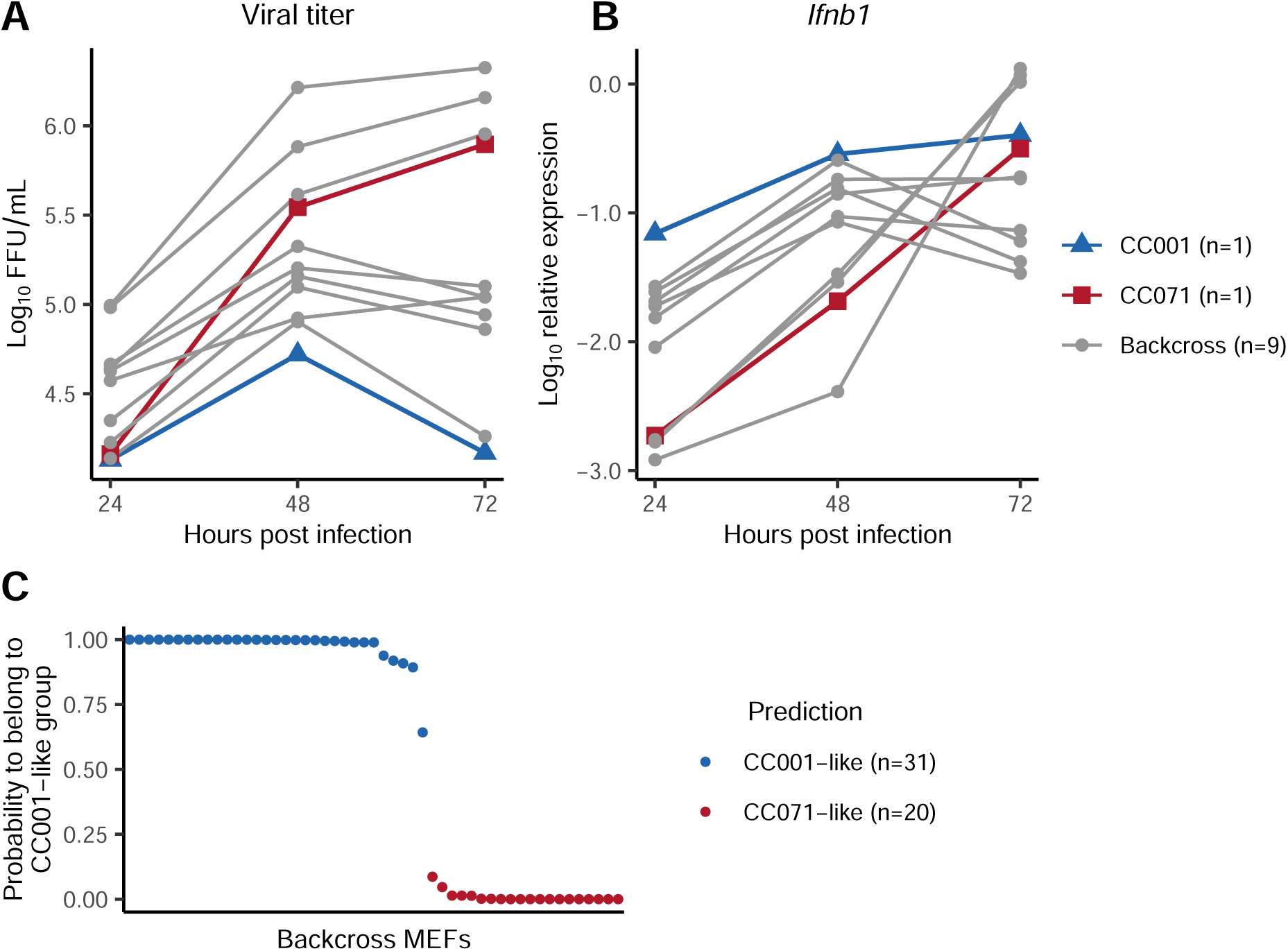
Backcross MEFs display either a CC001-like or a CC071-like phenotype. MEFs derived from CC001 (blue triangles), CC071 (red squares) and backcross (gray circles) embryos were infected with ZIKV at a MOI of 5. (A) Viral titer and (B) *Ifnb1* expression in ZIKV-infected MEFs from 9 backcross embryos. Experimental conditions were as in Figure 1A-B. Red and blue curves show the results for CC071 and CC001 MEFs, respectively, from the same infection experiment. (C) Results of LDA on the backcross MEFs. LDA coefficients were calculated from *Ifnb1* expression data in CC001 and CC071 infected MEFs, and applied to backcross MEFs. The graph shows the probability of each BC MEF to belong to the “CC001-like” group, resulting in two distinct populations shown in blue (”CC001-like”) and in red (”CC071-like”). n: number of BC MEFs assigned to each group.

**Figure 4.**
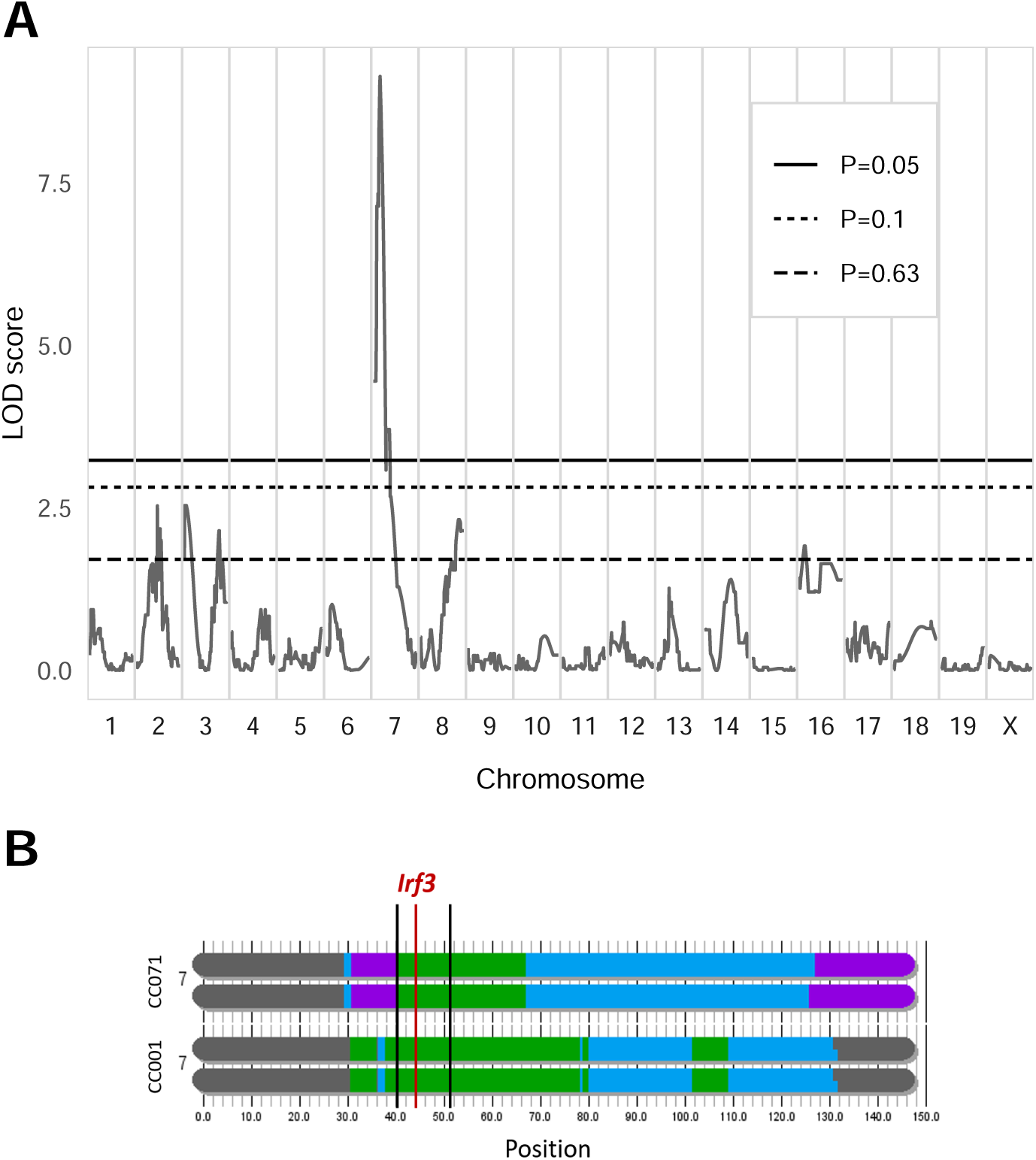
Genetic analysis of *Ifnb1* expression in backcross MEFs identifies a major determinant mapping to the *Irf3* locus. (A) Genome-wide linkage analysis of the LDA classification of backcross MEFs performed with R/qtl. X-axis: genomic location. Y-axis: LOD score of the phenotype-genotype association. Genome-wide significance thresholds (P = 0.05, P = 0.1 and P = 0.63) were computed from 1000 data permutation. The chromosome 7 peak has a LOD score of 9.138. (B) Schematic representation of CC001 and CC071 chromosome 7 haplotypes, from https://csbio.unc.edu/CCstatus/CCGenomes/. Colors represent the CC ancestral haplotypes (same colors as in Figure 2A). Thick vertical black lines show the peak’s 95% Bayesian confidence interval (25.9-31.3 cM, corresponding to 40.1-50.6 Mb). The red line shows the position of *Irf3* gene.

To confirm this apparently binary distribution, we conducted linear discriminant analysis (LDA) on CC001 and CC071 MEFs using *Ifnb1* expression at the three time points as variables. Applying the LDA coefficients to backcross MEFs data classified individuals either in a CC001-like group (n=31; 61%) or in a CC071-like group (n=20; 39%), with a mean probability of prediction of 0.975 and 0.991, respectively (Fig 3C). Quantitative trait locus (QTL) mapping was performed using LDA classification as a binary trait. Genome scan identified a peak on chromosome 7 with a LOD (logarithm of the odd) score of 9.138 (p < 0.001, Fig 4A) located in a region centered on the *Irf3* gene which, given its main role in the regulation of *Ifnb1* expression, appeared as an obvious candidate. However, both CC001 and CC071 inherited the CAST/EiJ haplotype in this region (Fig 4B), strongly suggesting that CC071’s susceptibility was caused by a variant proper to this strain.

### Abnormal splicing of Irf3 mRNA in CC071 leads to a loss of IRF3 transcriptional function

To identify the CC071-specific mutation, we re-analyzed the RNAseq data and investigated the splicing events between *Irf3* exons. As shown in Fig 5A, no splicing was observed between exons 6 and 7 in CC071 MEFs, while a short cryptic exon was added to exon 6 (red box). This aberrant splicing resulted in an mRNA lacking the last two exons. Notably, exon 8 encodes the serin-rich region of the protein with the phosphorylation sites necessary for IRF3 activation and nuclear translocation leading to *Ifnb1* transcription (Fig 5B). Neither long-range PCRs nor sequencing could identify the exact nature of *Irf3* genetic alteration in CC071 but suggested the insertion of a repeated sequence between exons 6 and 7. Nevertheless, the functional consequence of this mutation was confirmed by Western blot using a specific C-terminal IRF3 antibody which showed that full-length IRF3 protein was absent in CC071 MEFs (Fig 5C). Moreover, immunofluorescence using an antibody directed against phosphorylated IRF3 detected a positive signal in the nucleus of many ZIKV-infected CC001 and B6 MEFs, but not in CC071 MEFs (Fig 5D). Altogether, these results show that CC071 carries a mutation in *Irf3* that prevents IRF3 phosphorylation which is required to induce *Ifnb1* expression. Whether the altered mRNA sequence prevents the production of the protein or alters its activity, this mutation results in the loss of IRF3 transcriptional function.

**Figure 5.**
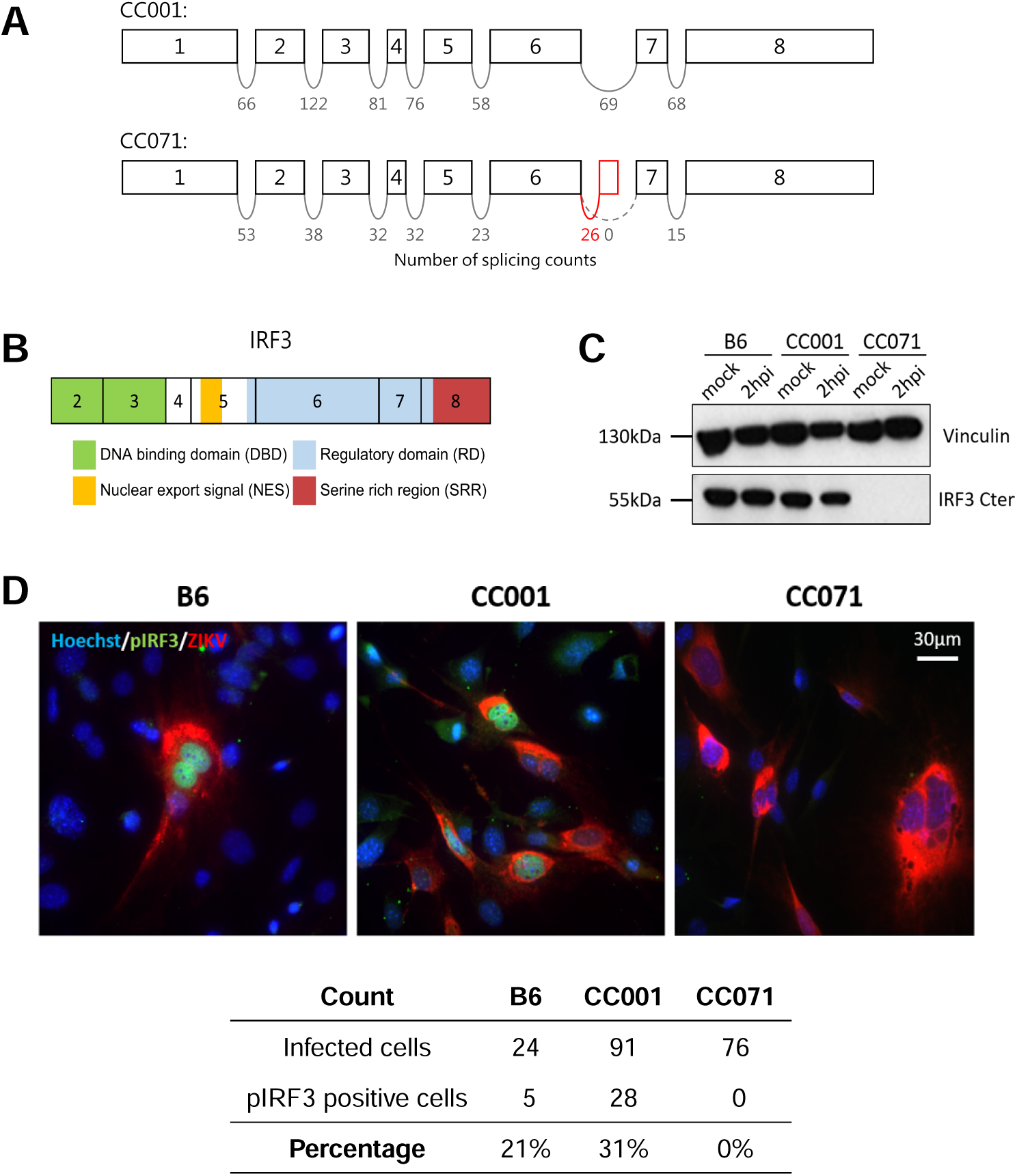
CC071 *Irf3* mRNAs show abnormal splicing, resulting in defective IRF3 protein. (A) Schematic representation of the exons of the *Irf3* gene with the number of reads spanning successive exons in the CC001 and CC071 RNAseq data (one sample of each strain). The red box between CC071’s exons 6 and 7 depicts a novel exon resulting from abnormal splicing. (B) Schematic representation of the IRF3 protein structural domains (exon 1 is untranslated). Exon 8 encodes the serine rich region containing the phosphorylation sites for IRF3 activation. (C) Western blot using an anti-C-terminal IRF3 antibody from mock-infected and ZIKV-infected B6, CC001 and CC071 MEFs at 2 hpi, showing the absence of full-length IRF3 in CC071 MEFs. Vinculin was used as a loading control. (D) Immunofluorescence using an anti-phosphorylated IRF3 (pIRF3, green) in ZIKV-infected B6, CC001 and CC071 MEFs at 24 hpi, showing the absence of pIRF3 in the nucleus of CC071 MEFs upon infection. Red-labeled 4G2 antibody labels ZIKV-infected cells. Cell nuclear DNA labeled by Hoechst (blue). Quantification of the number of infected and pIRF3 positive cells is presented in the table. Proportions were established on 420, 428 and 551 cells for CC001, CC071 and B6, respectively.

### CC071 Irf3 mutation is responsible for uncontrolled viral replication in MEFs

To test if the delayed *Ifnb1* expression resulting in uncontrolled viral replication in CC071 MEFs was caused exclusively by the *Irf3* mutation, we performed a quantitative complementation test by producing compound heterozygous MEFs carrying a knockout *Irf3* allele (*Irf3^KO^*) and a CC071 allele (*Irf3^71^*). These *Irf3^KO/71^* MEFs were compared with CC071 and B6-*Irf3^KO/KO^* MEFs, and with heterozygous MEFs carrying a B6 wildtype allele and either an *Irf3^KO^* or an *Irf3^71^* allele (*Irf3^+/KO^* or *Irf3^+/71^*, respectively, Fig 6A). While *Irf3^+/KO^* and *Irf3^+/71^* ZIKV-infected MEFs showed the same pattern as CC001 or B6 MEFs (rapid induction of *Ifnb1* expression and controlled viral replication, see Fig 1A for comparison), CC071 (*Irf3^71/71^*), B6-*Irf3^KO/KO^* and *Irf3^KO/71^* MEFs carrying two defective alleles at *Irf3* showed similar results (Fig 6B-C). These data demonstrate that, since the *Irf3^KO^*did not complement the *Irf3^71^* allele, the *Irf3* mutation in CC071 contributes to the defects observed in ZIKV-infected MEFs. Moreover, since the data obtained on CC071 and on *Irf3^KO/71^* MEFs were identical, we conclude that the CC071 *Irf3* mutation is sufficient to induce the defects observed in CC071 MEFs.

**Figure 6.**
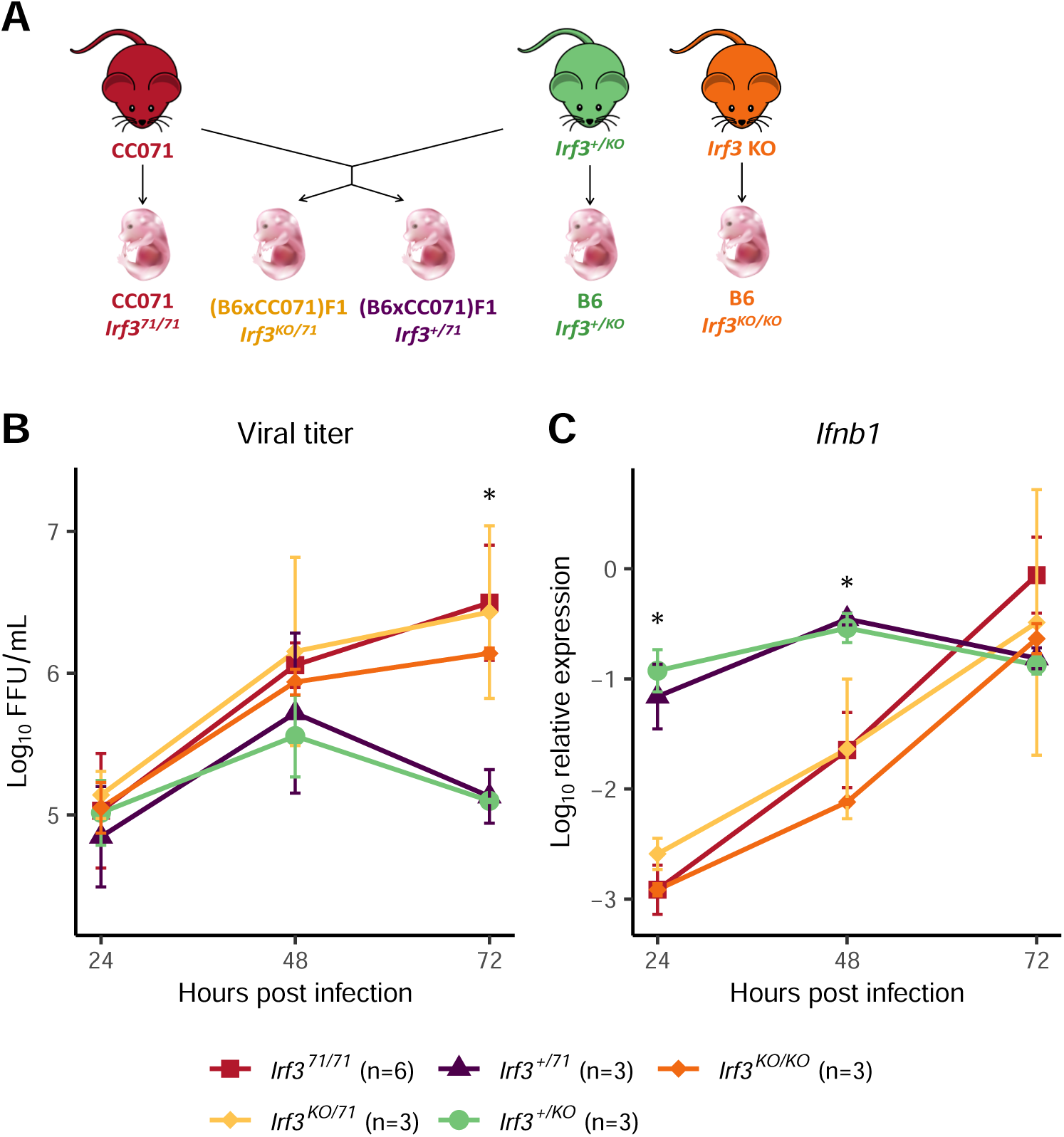
Quantitative complementation test confirms that *Irf3* LOF in CC071 is responsible for uncontrolled viral replication and delayed *Ifnb1* expression. (A) Mice heterozygous for an inactivated *Irf3* allele (B6-*Irf3^+/KO^*) were mated with CC071 mice to produce four types of embryos from which MEFs were derived. The genetic background and *Irf3* genotype is shown below each type of embryos. (B) Viral titer and (C) *Ifnb1* expression in ZIKV-infected MEFs. Experimental conditions were as in Figure 1A-B. Data are mean +/- sem from 6 biological replicates for CC071 and 3 for the other groups. The asterisks represent the results of ANOVA test between all groups (* p < 0.05). Results of the Tukey HSD post-hoc are as follows. Viral titer at 72h: CC071 vs *Irf3^+/71^* and vs *Irf3^+/KO^*: p < 0.001. *Irf3^KO/71^* vs *Irf3^+/71^* and vs *Irf3^+/KO^*: p < 0.01. *Irf3^KO/KO^* vs *Irf3^+/71^* and vs *Irf3^+/KO^*: p < 0.05. *Ifnb1* expression at 24hpi: CC071 vs *Irf3^+/71^* and vs *Irf3^+/KO^*, *Irf3^KO/71^* vs *Irf3^+/71^*and vs *Irf3^+/KO^*, *Irf3^KO/KO^* vs *Irf3^+/71^*and vs *Irf3^+/KO^*: p < 0.001. *Ifnb1* expression at 48hpi: CC071 vs *Irf3^+/71^*and vs *Irf3^+/KO^*, *Irf3^KO/71^*vs *Irf3^+/71^* and vs *Irf3^+/KO^*, *Irf3^KO/KO^*vs *Irf3^+/71^* and vs *Irf3^+/KO^*: p < 0.01.

### CC071 Irf3 mutation is not sufficient to explain susceptibility in vivo

We investigated whether this *Irf3* mutation was also responsible for the high susceptibility to ZIKV of CC071 mice. We first evaluated its effects in a context allowing IFN-I response. To this aim, we compared B6, B6-*Irf3* KO and CC071 mice infected without prior MAR1-5A3 treatment, which results in moderate and short-lasting plasma viral loads without clinical signs, as we previously reported (15). Similar viral loads were observed in B6 and B6-*Irf3* KO mice at all days p.i. (dpi), while CC071 mice showed significantly higher viral loads at days 1 and 2 p.i. (Fig 7A). This result indicates that, unlike in MEFs, the *Irf3* null mutation does not induce elevated viral replication *in vivo* (as measured by plasma viral load) and suggests that CC071 mice carry susceptibility alleles at other loci.

**Figure 7.**
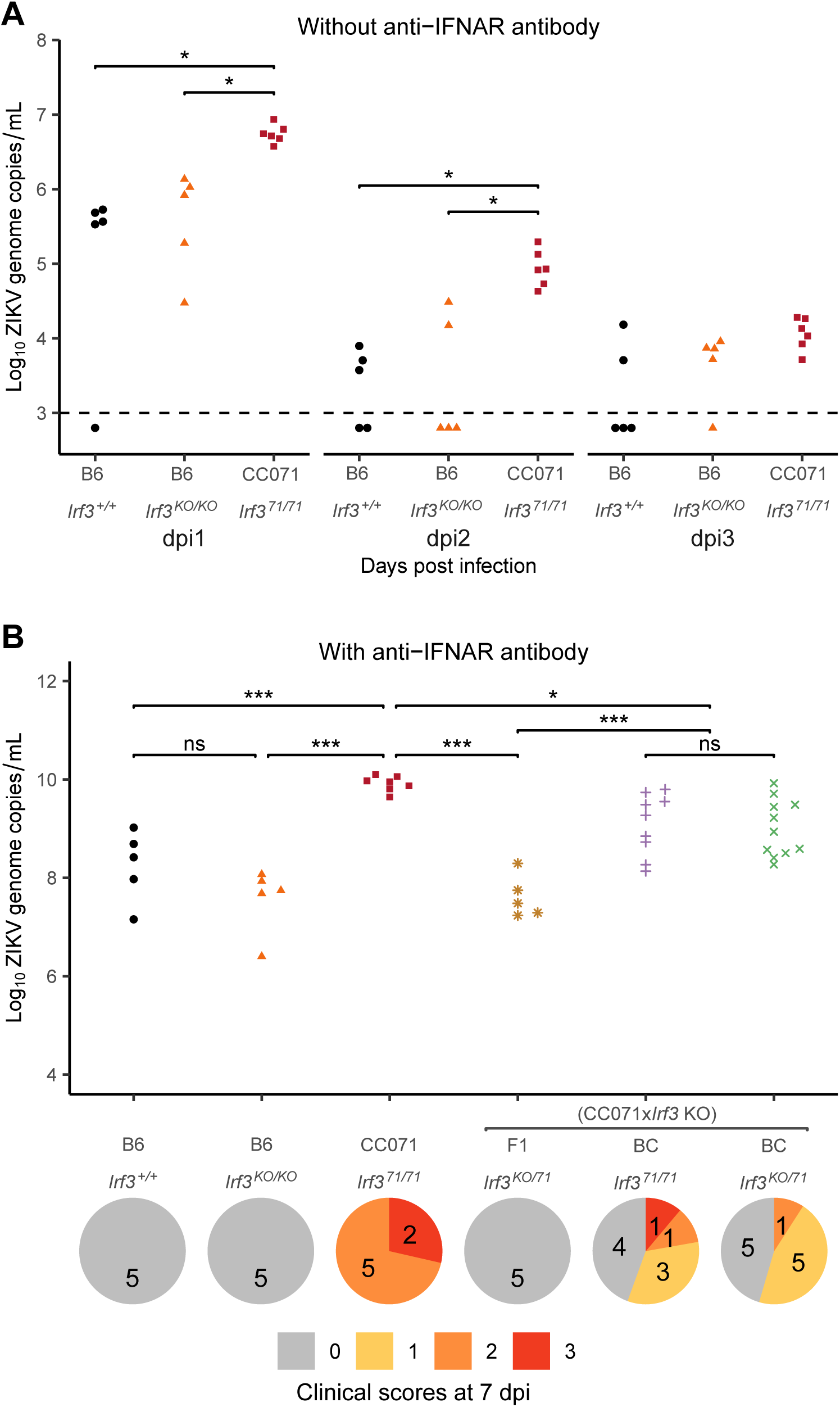
*Irf3* LOF is not sufficient to explain CC071 susceptibility to ZIKV infection *in vivo*. (A) B6, B6-*Irf3* KO and CC071 mice were infected IP with 10^7^ FFUs of ZIKV and monitored for 7 days, without prior IP injection of MAR1-5A3 IFNAR-blocking monoclonal antibody. Plasma viral load was quantified at days 1 to 3 p.i. by RT-qPCR. The two lines of the group legend (X-axis) indicate the genetic background of the mice and their genotype at the *Irf3* locus, respectively. Each dot represents one mouse. Groups were compared by Kruskal-Wallis followed by Wilcoxon test with Holm correction for multiple testing (* p < 0.05). (B) B6, B6-*Irf3* KO, CC071, (CC071 x *Irf3* KO) F1 and (CC071 x *Irf3* KO) x CC071 BC mice were infected IP with 10^7^ FFUs of ZIKV after IP injection of 2 mg of MAR1-5A3 IFNAR-blocking monoclonal antibody 24 h before infection, and monitored for 7 days. The graph shows plasma viral loads quantified at day 2 p.i. by RT-qPCR. The two lines of the group legend (X-axis) indicate the genetic background of the mice and their genotype at the *Irf3* locus, respectively. BC mice are separated into two groups depending on the genotype at the *Irf3* gene: homozygous for the CC071 mutant allele (*Irf3^71/71^*), or heterozygous for the CC071 and the KO alleles (*Irf3^KO/71^*). *Irf3* genotyping results are presented in Figure S5. Each dot represents one mouse. Groups were compared by ANOVA followed by Tukey HSD (* p < 0.05, ** p < 0.01, *** p < 0.001). Below the graph is shown the distribution of clinical scores at day 7 p.i. in the same groups of mice as above (0, no symptoms; 1, slight hunched posture; 2, ruffled fur, hunched posture and/or mild ataxia; 3: prostration, ataxia, partial paralysis).

We then blocked IFN-I response by pre-treating mice with MAR1-5A3 and compared peak plasma viral load and clinical signs in ZIKV-infected B6, B6-*Irf3* KO and CC071 mice. While no differences were observed between B6 and B6-*Irf3* KO mice (moderate plasma viral loads without clinical signs), as expected if the IFN-I response is neutralized, CC071 mice developed significantly higher viral loads at day 2 p.i. and signs of disease (ruffled fur, hunched posture and body weight loss) around 7 days p.i. (Fig 7B). This contrast could result either from functional differences between the *Irf3^KO^* and *Irf3^71^* alleles or from CC071 alleles at other genes not linked with the IFN-I response. F1 mice between CC071 and B6-*Irf3* KO mice gave similar results as B6-*Irf3* KO mice, suggesting that CC071 susceptibility alleles were recessive. Therefore, we generated a small cohort of (CC071 x B6-*Irf3* KO) x CC071 backcross mice and genotyped them for the *Irf3* gene by PCR using primers that amplify intron 6 of the *Irf3^KO^* allele, but not of the *Irf3^71^*allele. (Fig 5A and Fig S5). We found no differences in plasma viral load and clinical scores between *Irf3^KO/71^* and *Irf3^71/71^* backcross mice, indicating that the two alleles are functionally equivalent *in vivo*. Moreover, *Irf3^KO/71^* backcross mice were more severely affected than *Irf3^KO/71^* F1 mice, indicating the strong effects of recessive alleles from CC071 which likely explain the variability of plasma viral load and clinical signs among backcross mice. Altogether, our results demonstrate that the CC071 mutation is a loss-of-function (LOF) variant which does not confer susceptibility to ZIKV disease in the absence of other recessive alleles carried by CC071 at genes not associated with the IFN-I response.

## Discussion

An increasing number of publications have reported how the genetic diversity provided by the CC translates into an extended range of resistant and susceptible phenotypes in various infectious models. Since our first description of the high susceptibility of CC071 mice to ZIKV infection, with disease severity and peak plasma viral load almost as high as those of B6-*Ifnar1* KO mice, and higher clinical signs and mortality than 129-*Ifnar1* KO mice (15), several groups have reported similar observations for other viruses. High viral load was found in CC071 mice after dengue infection (15). Lethality after infection with West Nile virus (15) or Powassan virus (16) were also described, although there is likely contribution from the defective *Oas1b* allele that CC071 has inherited from 129S1/SvImJ. CC071 was also one of the most susceptible CC strains to Rift Valley Fever virus infection (17) and to hepacivirus with long-term viral persistence (21). Because of its unique genetic background, this strain will be increasingly useful for infectious diseases studies, which underlines the importance of deciphering the mechanisms of its susceptibility to each pathogen.

In this study, we leveraged from our previous observation that CC071 MEFs failed to control ZIKV replication by comparison with resistant strains. Although they cannot recapitulate the complex interactions between multiple pathways, cell types and tissues of a whole organism, MEFs are a convenient cellular model which can be easily derived from any mouse genetic background. We identified a delayed activation of the *Ifnb1* gene in CC071 MEFs, resulting in delayed stimulation of ISGs. We then used a combination of genetic approaches to find the causative gene defect. The observation that MEFs were normally responsive to IFN-I stimulation was consistent with the higher susceptibility of mice treated by the MAR1-5A3 antibody compared with untreated mice which did not develop symptoms and showed much lower plasma viral load (15). These results pointed at the pathway from PAMP sensors to *Ifnb1* transcription factors. Transcriptional analysis did not identify reduced expression of non-ISGs of this pathway. CC strains’ genomic structure allows searching for haplotypes inherited from the same founder in CC strains showing similar phenotypes. However, we did not identify such haplotypes for genes of the *Ifnb1* induction pathway, suggesting that the defects observed in CC071 MEFs were strain specific. It is finally the analysis of MEFs derived from backcross embryos that established a monogenic inheritance, and genetic linkage unambiguously pointed at the causative *Irf3* gene. The MEF experimental model was particularly appropriate since we could derive cell lines from every backcross embryo. Our RNAseq data showed an RNA splicing defect in the *Irf3* gene in CC071, which functional consequences could be validated *in vitro*. The formal proof that the CC071 *Irf3* mutation was necessary and sufficient to cause *Ifnb1* delayed activation and uncontrolled viral replication came from a quantitative complementation test in which the *Irf3^KO^* allele was combined either with a CC071 or a B6 (functional) allele. Altogether, these results show that the *Irf3* defect in CC071 abrogates its transcriptional activity. Unfortunately, none of the antibodies recognizing the N-terminal protein domain that we tested worked in our hands in western blot. Therefore, we could not establish whether the *Irf3* mRNA detected by RNAseq is translated into an abnormal protein or not. However, the *in vivo* observation that *Irf3^KO/71^* and *Irf3^71/71^*backcross mice displayed the same phenotypes upon ZIKV infection shows that the two alleles are functionally equivalent in this context.

*Irf3* maps to a chromosomal region that CC071 inherited from CAST/EiJ, like the resistant CC001 strain (and CC046/Unc, not available to us). The CC071-specific mutation may have arisen during the inbreeding generations leading to this strain. Alternatively, the mutation might have been segregating in the CAST/EiJ founders and transmitted by chance only to the CC071 breeding funnel. Since all CAST/EiJ founders used in CC crosses originated from The Jackson Laboratory colony, genotyping past and present breeders of this colony could not only unravel the origin of the mutation but also allow eliminating this mutation to avoid unwanted effects.

Notably, this is not the first example of a CC-strain-specific mutation. We (22) and others (23) previously reported the extreme susceptibility of CC042/GeniUnc (CC042) to *Salmonella* Typhimurium and to *Mycobacterium tuberculosis*, respectively, as a consequence of a *de novo* 15-nucleotide deletion in the *Itgal1* gene. In the case of *Salmonella* Typhimurium, CC042 was standing out, with bacterial loads up to 1000 times higher than other CC and the susceptible B6 strains. In our previous study on ZIKV, CC071 was the most susceptible strain, but its peak viral load was just the highest in a continuous distribution of values. Additionally, this is not the first example of a *Irf3* spontaneous variant identified in mice modulating susceptibility to bacterial (24) or viral infections (25).

*Irf3* is an important transcription factor involved in the innate immune response. It is constitutively expressed and, at rest, inactive IRF3 is present in the cytoplasm. Upon viral entry (or other stimuli that activate TLRs such as TLR3 and 4 or RIG-I-like receptors), signal transduction leads to the phosphorylation of IRF3, resulting in its dimerization and translocation to the nucleus where it binds to the *Ifnb1* promoter (26). This mechanism leads to the very fast production of IFNβ which is secreted by the cell and triggers the immediate response to viral infection through the activation of ISGs with diverse antiviral functions (27). In addition to its transcriptional activity, IRF3 has been shown to counter viral infection by two additional mechanisms. IRF3 induces apoptosis through the RLR-induced IRF3-mediated pathway of apoptosis (RIPA) in infected cells (28) and inhibits NF-κB-mediated inflammation through the repression of IRF3-mediated NF-κB activity (RIKA) (29). These newly described roles of IRF3 depend on other protein domains than those involved in its transcriptional activity. Whether these activities are also abrogated by the CC071 *Irf3* mutation remains to be established.

In mice, studies that used *Irf3*-deficient cellular models have consistently reported decreased IFN-I production and/or increased viral replication. For example, after infection with WNV, viral replication was increased in B6-*Irf3* KO bone marrow macrophages (BMMs) and moderately in primary neurons (30). Higher viral replication was observed in herpes simplex virus 1 (HSV-1)-infected B6-*Irf3* KO bone marrow-derived dendritic cells (BMDCs) and BMMs, and IFNβ was reduced in BMDCs supernatants (31). Similarly, *Ifnb1* expression was reduced in HSV-1-infected (32) and CHIKV-infected (33) B6-*Irf3* KO MEFs. In line with these findings, our study provides, to our knowledge, the first evidence for a role of *Irf3* in the infection of murine cells by ZIKV. Moreover, we recently reported that primary cultured neurons (PCNs) show a delayed activation of the *Ifnb1* expression upon ZIKV exposure compared with MEFs, and that this delay is even longer in PCNs derived from CC071 compared with CC001 (34). The identification of the *Irf3* mutation in CC071 provides an explanation for this observation, although it would require confirmation in B6-*Irf3* KO PCNs.

In contrast, *in vivo* studies have reported inconsistent consequences of *Irf3* deficiency between viral infections. Here, we did not observe any differences in clinical signs nor in plasma viral loads between B6 and B6-*Irf3* KO mice (pre-treated or not with the MAR1-5A3 antibody), consistently with a previous study that reported neither mortality, nor body weight loss after ZIKV infection (10). Likewise, B6-*Irf3* KO mice have been reported to show no mortality and low virus load in the circulation following dengue virus (35) and CHIKV infection (33). Contrastingly, WNV infection was lethal in all B6-*Irf3* KO mice while 65% of infected WT mice survived (30). Differences were also associated with the route of infection. For example, all WT and B6-*Irf3* KO mice survived after intravenous inoculation with HSV-1 (32), while intranasal inoculation led to 90% mortality in B6-*Irf3* KO but only 30% in WT mice (36).In humans, LOF mutations in *Irf3* have been associated with increased susceptibility to diseases caused by WNV (37,38), HSV-1 (39,40) and more recently by SARS-CoV-2 (41). Discrepancies between the consequences of *Irf3* deficiency in *in vitro* and *in vivo* experiments likely reflect differences between a single cell type model with an intrinsic capacity to mount a more or less IRF3-dependent IFN-I response, and a multicellular organism in which the paracrine effect of IFN-I produced by the most reactive cells induces protection of other, less reactive, cell types. In fact, we have recently reported such cooperation between neurons and microglia cells (34). Our *in vivo* data, showing high plasma viral loads and clinical signs in all CC071 and half of backcross mice after MAR1-5A3 treatment, demonstrated that the high susceptibility of CC071 mice requires the contribution of other recessive alleles. Since their effect was observed in mice which IFNAR receptor had been blocked, we conclude that their mode of action is not dependent on an intact IFN-I response. Moreover, the comparison between the results of the complementation test on MEFs, with CC071 and *Irf3^KO/71^* yielding very similar data, and the *in vivo* observation that CC071 showed higher viremia than *Irf3^KO/71^* F1 mice, suggests that these other susceptibility alleles may not act at the cell level, at least during the first 72 hpi, but rather at a more integrated or systemic level. Whether or not these alleles require *Irf3* deficiency to induce the susceptibility observed in the CC071 mice has yet to be determined. While our limited understanding of the severe disease developing in CC071 ZIKV-infected mice does not point at potential mechanisms, identifying these alleles will allow disentangling the genetic factors controlling ZIKV and other viruses’ pathogenesis.

## Material and methods

### Mice and crosses

C57BL/6J (B6) mice were purchased from Charles River Laboratories France. Collaborative Cross strains (CC001/Unc, CC071/TauUnc, CC005/TauUnc, CC011/Unc, CC026/GeniUnc, CC061/GeniUnc, CC021/Unc, CC006/TauUnc, CC025/GeniUnc, CC039/Unc, CC060/Unc) were purchased from the Systems Genetics Core Facility, University of North Carolina and bred at the Institut Pasteur. *Irf3 Irf7* double KO mice (C57BL/6J-*Bcl2l12*/*Irf3^tm1Ttg^ Irf7^tm1Ttg^*,(32,42)) were bred at the Institut Pasteur and backcrossed to B6 mice to generate *Irf3* single KO mice (B6-*Irf3* KO). Genetic mapping was performed on MEFs derived from (CC001 x CC071) x CC071 and CC071 x (CC001 x CC071) backcross embryos. For the quantitative complementation test, MEFs were derived from CC071, B6-*Irf3* KO and (B6-*Irf3^+/KO^* x CC071) embryos.. *In vivo* experiments were performed on B6, B6-*Irf3* KO, CC071, (CC071 x *Irf3* KO) F1 and CC071 x (CC071 x *Irf3* KO)backcross mice. All crosses are described as female x male. All mice were maintained as described previously (15). Mouse experiments were approved by the Institut Pasteur Ethics Committee and authorized by the French Ministry of Research (project #19469), in compliance with French and European regulations.

### ZIKA virus

The FG15 Asian Zika virus (ZIKV) strain, isolated from a patient during ZIKV outbreak in French Guiana in December 2015, was obtained from the Virology Laboratory of the Institut Pasteur of French Guiana. Viral stock (passage 5) was prepared from supernatant of infected C6/36 cells, clarified by centrifugation at 800g and titrated on Vero cells by focus-forming assay.

### Mouse infection

All infection experiments were performed in a biosafety level 3 animal facility and mice were kept in isolators. Six- to 10-week-old, male or female mice were injected intraperitoneally with 10^7^ PFU of ZIKV FG15. In some *in vivo* experiments, mice received an IP injection of 2 mg of MAR1-5A3 anti-IFNAR antibody (Euromedex, Cat#BX-BE0241) one day prior infection. Mouse numbers are indicated in figure legends. Both males and females were used since no differences between sexes were detected in our previous and present experiments. Clinical signs and body weight loss were recorded for up to seven days post infection. Blood samples were collected on EDTA from the retromandibular vein for plasma viral load assessment. Quantification of ZIKV viral copies by qPCR was previously described (15).

### MEFs isolation

Pregnant females were euthanized at day 13.5-15.5 of gestation. For B6, CC001, CC071 and crosses used for genetic mapping and the complementation test, MEFs were isolated from individual fetuses to obtain biological replicates. For other CC strains, MEFs were derived from individual or pooled fetuses. Fetus bodies were chopped and digested with trypsin (Gibco Cat#25300054), then cultured at 37°C and 5% CO_2_ in complete medium (DMEM Gibco Cat# 31966047, 10% fetal bovine serum PAA Laboratories Cat#A15-101, 1% penicillin/streptomycin Sigma Cat#P4333). MEFs were used until passage 2. For the backcross experiment, MEF lines were isolated from 51 backcross fetuses. Heads were used to prepare DNA for whole-genome genotyping.

### MEFs infection

MEFs were seeded at 5.10^4^ cells per well in 24-well plates the day before infection. They were exposed to ZIKV FG15 strain at a MOI of 5 for 2 hours after which the inoculum was replaced with fresh complete medium and MEFs were incubated for up to 72 hours. For kinetics studies, different wells were used for each time point. Backcross MEFs were infected in 6 infection experiments, each of which included one CC001 and one CC071 MEF lines.

### MEFs IFNα stimulation

MEFs were seeded at 5.10^4^ cells per well in 24-well plates one day before stimulation and treated with 300 IU/mL IFNα (Miltenyi Biotec Cat#130-093-131) and incubated for up to 24 hours.

### Focus forming assay

Quantification of ZIKV particles was performed by focus forming assay on Vero cells (ATCC CRL-1586) as previously described (15).

### RNA extraction from cells

MEFs were lysed in 350µL of RLT buffer (Qiagen) with 1% β-mercaptoethanol. RNA was extracted using RNeasy Mini Kit (Qiagen Cat#74104) according to the manufacturer’s instructions, with addition of DNase I (Qiagen Cat# 79254) to prevent genomic DNA contamination.

### Reverse transcription and qPCR

Reverse-transcription was performed on 200ng of RNA using Superscript II polymerase (Invitrogen Cat#18064022) and RNaseOUT ribonuclease inhibitor (Invitrogen Cat#10777019). qPCR was performed on 20ng of cDNAs using Power SYBR Green PCR Master Mix (Applied Biosystems Cat#4367659) and 6pmol of each primer, on a QuantStudio 12K Flex or a ViiA 7 (ThermoFisher Scientific). Primers (Eurofins) sequences are provided in Sup Table 1. Gene expression was expressed on a Log10 scale of relative expression to the reference *Tbp* gene.

### Genotyping

Genomic DNA was prepared from backcross fetuses’ heads by proteinase K digestion, phenol-chloroform extraction and ethanol precipitation according to standard protocols. Whole-genome genotyping was performed at Neogen (Neogen/Geneseek, Inc, Lincoln, NE, USA) using the MiniMUGA array containing 11,125 SNP markers. For the quantitative complementation test, *Irf3* and *Irf7* genotyping was performed by Transnetyx (Cordova, TN) by real-time PCR on fetuses’ heads. (CC071 x *Irf3* KO) x CC071 backcross individuals were genotyped by PCR using primers amplifying the intron 6 of *Irf3^+^* and *Irf3^KO^* but not *Irf3^71^* alleles. Primers (Eurofins) sequences are provided in Sup Table 1.

### RNA splicing analysis from RNA sequencing data

RNA sequencing data were produced as described in the Supplementary methods. Splicing analysis of the *Irf3* gene was performed using Majiq 2.4 (43) with default parameters to investigate alternative transcripts between genotypes.

### Immunofluorescence

MEFs were plated on glass coverslips before infection, fixed with 4% paraformaldehyde for 20 min and permeabilized with 100% methanol for 10 min at -20°C. Cells were incubated with blocking buffer (5% FBS 0.3% triton in PBS) for 1 hour, with primary antibodies diluted in antibody incubation buffer (AIB: 1% BSA 0.3% triton in PBS) overnight at 4°C and with secondary antibodies and Hoechst (dilution 1:1000) diluted in AIB for 1 hour. Coverslips were mounted on slides and imaged with a widefield microscope (Zeiss Axio Observer.Z1 with a Plan-Apochromat 20x/0.8 M27 objective and a Hamamatsu sCMOS ORCA-Flash 4.0 v3 camera). ZEN blue 2012 software (ZEISS) imaging software was used for image capture and Image J software (National Institutes of Health) to adjust brightness and contrast. Primary and secondary antibodies are indicated in Sup Table 2.

### Western blot

MEFs were trypsinized for 5 min, washed and lysed in a protein extraction buffer (10mM TrisHCl pH7.5, 5mM EDTA, 150mM NaCl, 30mM Na_2_HPO_4_, 50mM NaF, 10% glycerol, 1% NP40, 1X cOmplete (Roche #11873580001), 1X PhosSTOP (Roche #4906845001), 1/1000 benzonaze (Sigma Cat#E1014)) for 30 min at 4°C. Proteins diluted in Laemmli were resolved on 4-12% Bis-Tris gels (Invitrogen Cat#NP0323BOX) in MOPS buffer (Invitrogen Cat#NP0001) and transferred to nitrocellulose membranes (Bio-Rad Cat#1620112) in a 25mM Tris 200mM glycine 20% ethanol buffer. Blots were blocked in 5% milk in TBS-T (0.1% Tween20 in Tris Base Sodium), incubated with primary antibodies diluted in 3% milk in TBST overnight at 4°C, and incubated with secondary antibodies diluted in 3% milk in TBST for 90 min. Blots were revealed with ECL substrate (Thermo Scientific Cat#32132) and imaged with X-ray films. Primary and secondary antibodies are indicated in Sup Table 2.

### Statistical analysis

Statistical analyses were performed using R version 4.1.0 (44). Viral titers, gene expression and genome copies were log-transformed for graphs and statistical tests. One way ANOVA followed by Tukey HSD were used for testing multiple comparisons. For *in vivo* studies without MAR1-5A3 treatment, non-parametric Kruskal-Wallis followed by Wilcoxon tests with Holm correction for multiple testing were used to handle values below the limit of detection. Linear discriminant analysis (LDA) was conducted using the MASS package (45). The LDA was trained on the phenotypes of the two parental CC001 and CC071 strains from each infection batch. LDA coefficients were applied to backcross mice for assignment to “CC071-like” or “CC001-like” groups.

### Genetic analysis

Raw genotypes were curated using the stuart package (46). QTL mapping was performed using R/qtl (47). LDA prediction was used as a binary trait. Statistical significance thresholds were computed by data permutation (n=1000). 95% confidence interval was estimated using the Bayesian method.

## Acknowledgements

We thank Matthieu Prot, Maxime Chazal and Sandrine Vandormael-Pournin for technical advice and for providing reagents. We are grateful to Tommy Penel and Rachid Chennouf of the Institut Pasteur Central Animal Facility of the C2RA (Center for Animal Resources and Research) for the careful breeding of CC strains and for the maintenance of mice in the BSL-3 animal facility, respectively. We thank Etienne Simon-Loriere and Nolwenn Jouvenet for scientific advice and continuous support along this project.

## Author contributions

X.M. conceived and supervised this study. M.B. and C.M. designed and performed the experiments and analyzed data. L.C. and C.R.P. performed experiments. E.K. analyzed RNAseq data. X.M. designed experiments and analyzed data. E.B. provided conceptual advice. M.B., C.M. and X.M. wrote the paper. All authors commented on and edited the manuscript.

## Funding

This work was supported by grants from the Agence Nationale de la Recherche (ANR) NeuroZika (ANR-20-CE16-0017) and from the French Government’s Investissement d’Avenir program, Laboratoire d’Excellence: IBEID (Integrative Biology of Emerging Infectious Diseases, ANR-10-LABX-62-IBEID). C.M. and M.B. were supported by doctoral fellowships from grant ANR-10-LABX-62-IBEID.

## Availability of data and materials

RNAseq primary data are available from the European Nucleotide Archive (accession number E-MTAB-12765). All other experimental data that support the finding of this study are available from the corresponding authors upon reasonable request.

## Competing interests

The authors declare that they have no competing interests.

## Legend to supplemental figures

**Figure S1.**
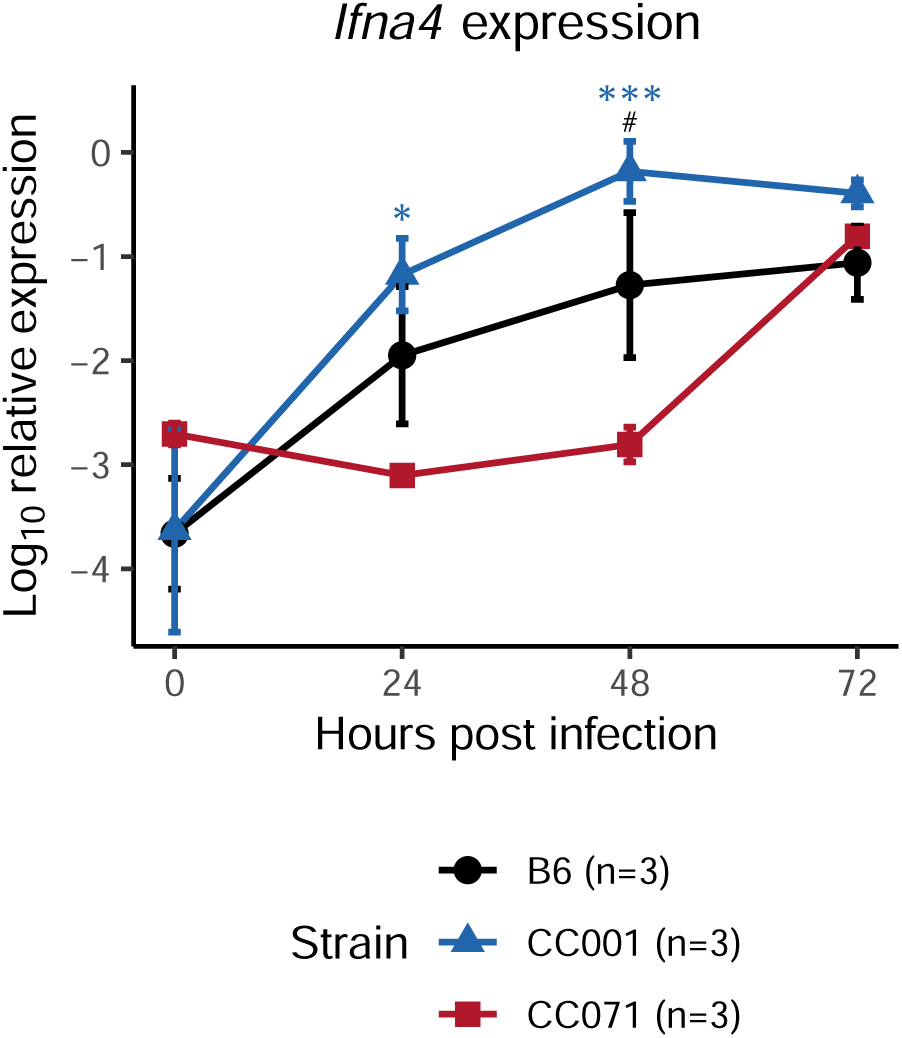
*Ifna4* expression is delayed in ZIKV-infected CC071 MEFs. MEFs derived from B6 (gray circles), CC001 (blue triangles) and CC071 (red squares) were infected with ZIKV at a MOI of 5. *Ifna4* expression was determined by RT-qPCR on MEFs total RNA by normalizing to *Tbp* housekeeping gene. Data are mean +/- sem from 3 biological replicates. For one CC001, one B6 and one CC071 replicates at 0 hpi and one CC071 replicate at 24 hpi, gene expression was below the limit of detection. Blue asterisks and black hashes show statistical significance of CC071 compared to CC001 and to B6, respectively (ANOVA followed by post-hoc Tukey HSD, */# p < 0.05, *** p < 0.001).

**Figure S2.**
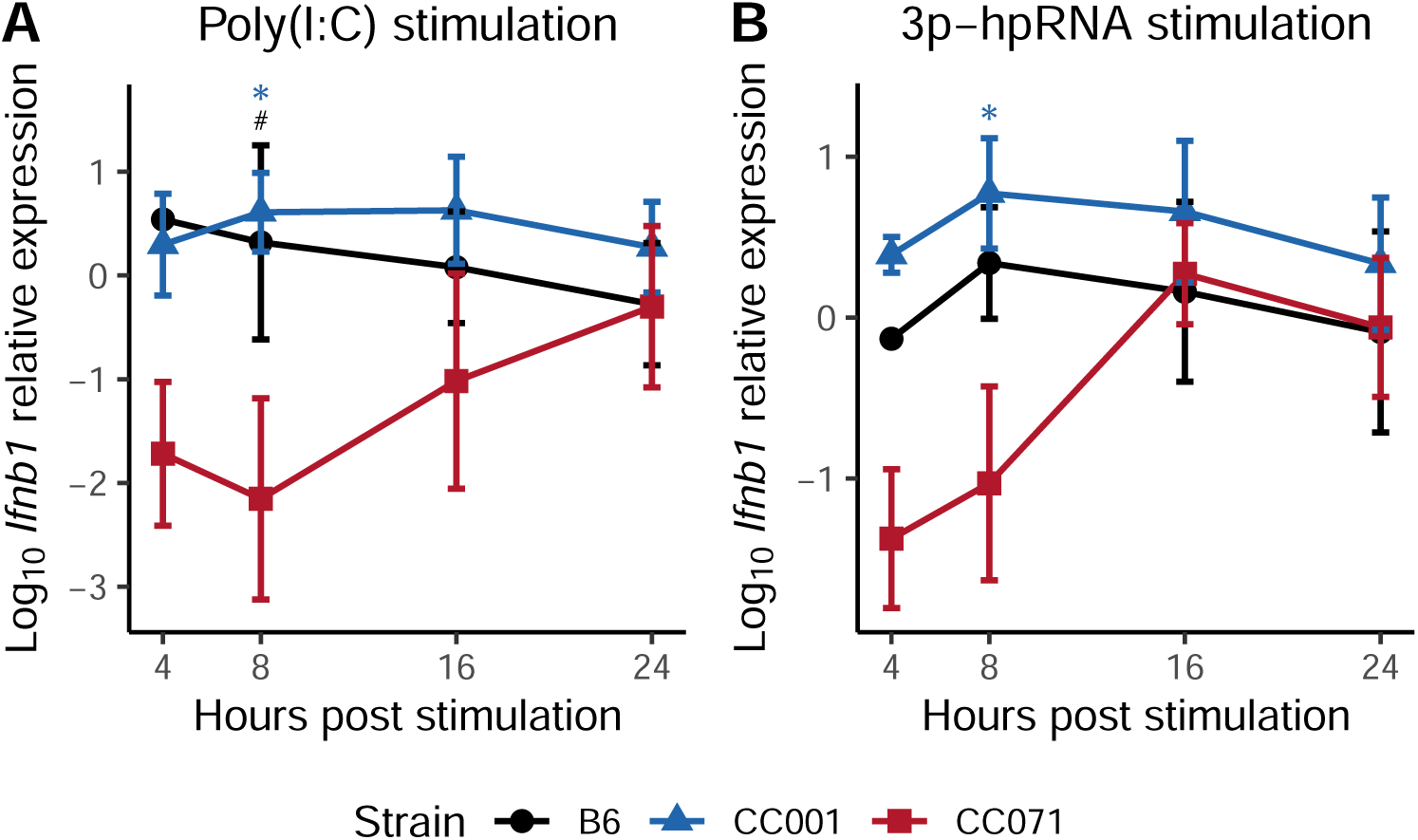
Delayed IFN-I expression in CC071 MEFs results from a constitutive defect in the transcription activation cascade. MEFs were transfected with either (A) poly(I:C), which activates both TLR and RIG-I pathways, or (B) 3p-hpRNA, a RIG-I agonist. *Ifnb1* expression was determined as in Figure 1B. Data are mean +/- sem from 3 biological replicates for CC001 and CC071 (2 at 4 hours) or 2 biological replicates for B6 (1 at 4 hours). Blue asterisks and black hashes show statistical significance of CC071 compared to CC001 and to B6, respectively (ANOVA followed by post-hoc Tukey HSD, */# p < 0.05).

**Figure S3.**
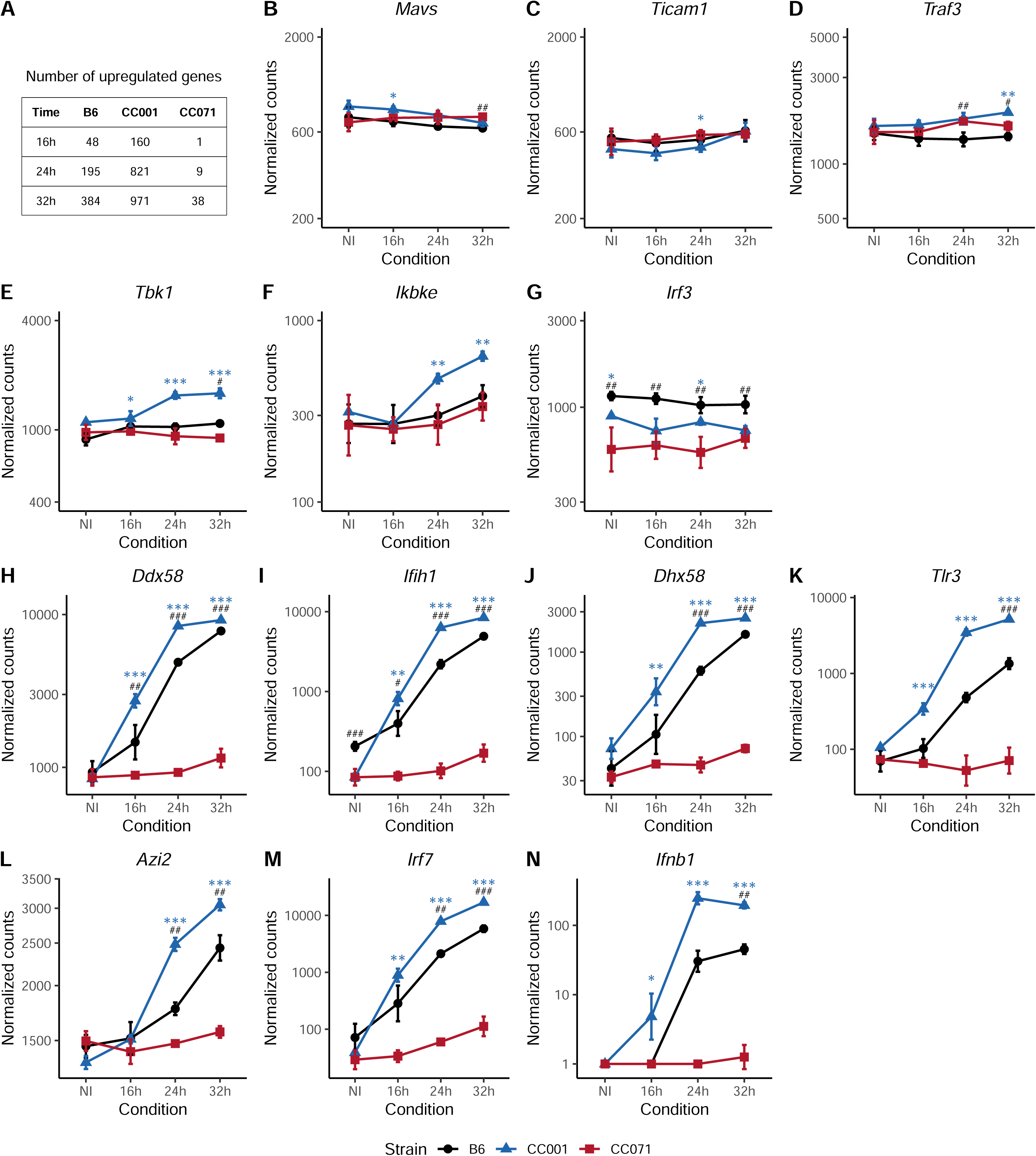
Expression of genes involved in the *Ifnb1* induction pathway in ZIKV-infected MEFs of B6, CC001 and CC071 strains. MEFs derived from B6 (black circles), CC001 (blue triangles) and CC071 (red squares) were infected with ZIKV at a MOI of 5. mRNA expression levels were measured by RNAseq in non-infected (NI) and ZIKV-infected MEFs. (A) Number of upregulated genes per strain at 16, 24 and 32 hpi (log2 fold-change > 1, FDR = 0.05). (B-G) Genes constitutively expressed. (H-N) Genes which expression is induced by the IFN-I response (ISGs). Expression levels are shown on a logarithmic scale. For *Ifnb1* expression, null counts were transformed to 1. Data are mean +/- sem from 3 biological replicates. Blue asterisks and black hashes show statistical significance of CC071 compared to CC001 and to B6, respectively (ANOVA followed by post-hoc Tukey HSD, */# p < 0.05, **/## p < 0.01, ***/### p < 0.001).

**Figure S4.**
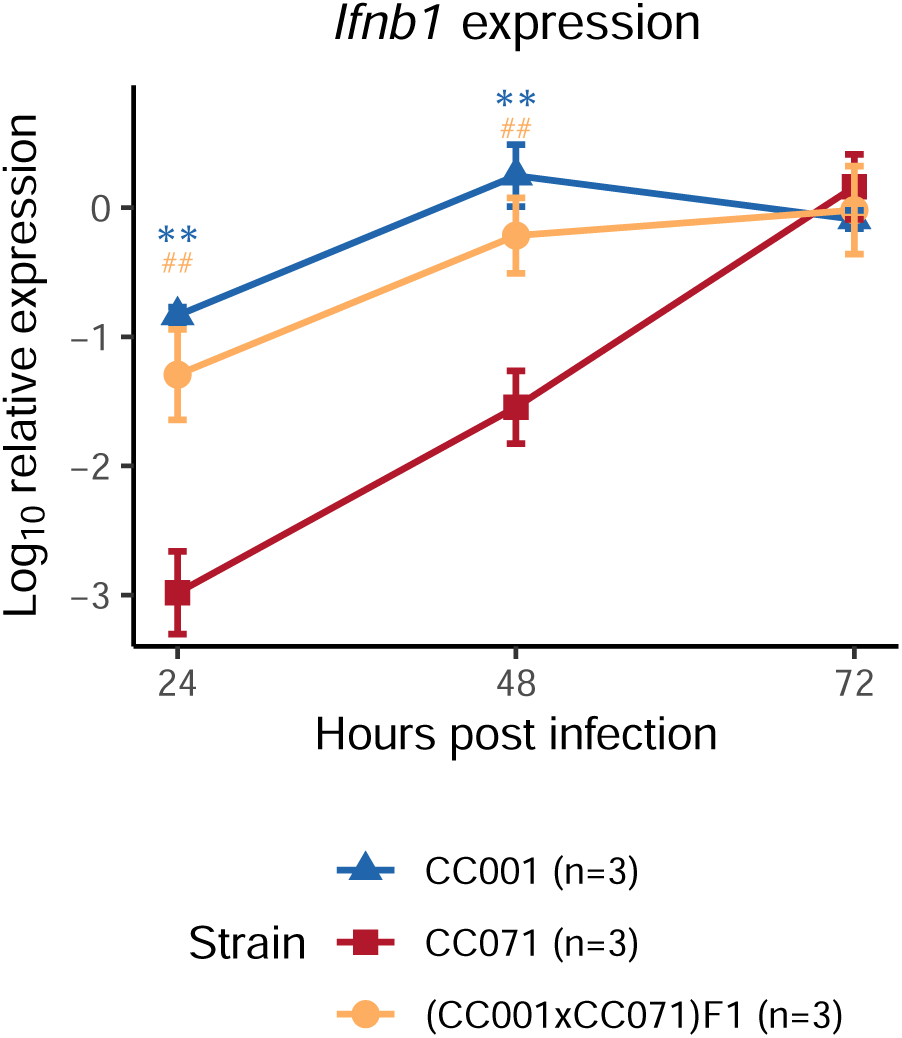
(CC001xCC071)F1 MEFs show normal induction of *Ifnb1*. *Ifnb1* expression upon ZIKV infection in CC001, CC071 and (CC001xCC071)F1 MEFs determined as in Figure 1B. Data are mean +/- sem from 3 biological replicates. Blue asterisks and orange hashes show statistical significance of CC071 compared to CC001 and to F1, respectively (ANOVA followed by post-hoc Tukey HSD, **/## p < 0.01).

**Figure S5.**
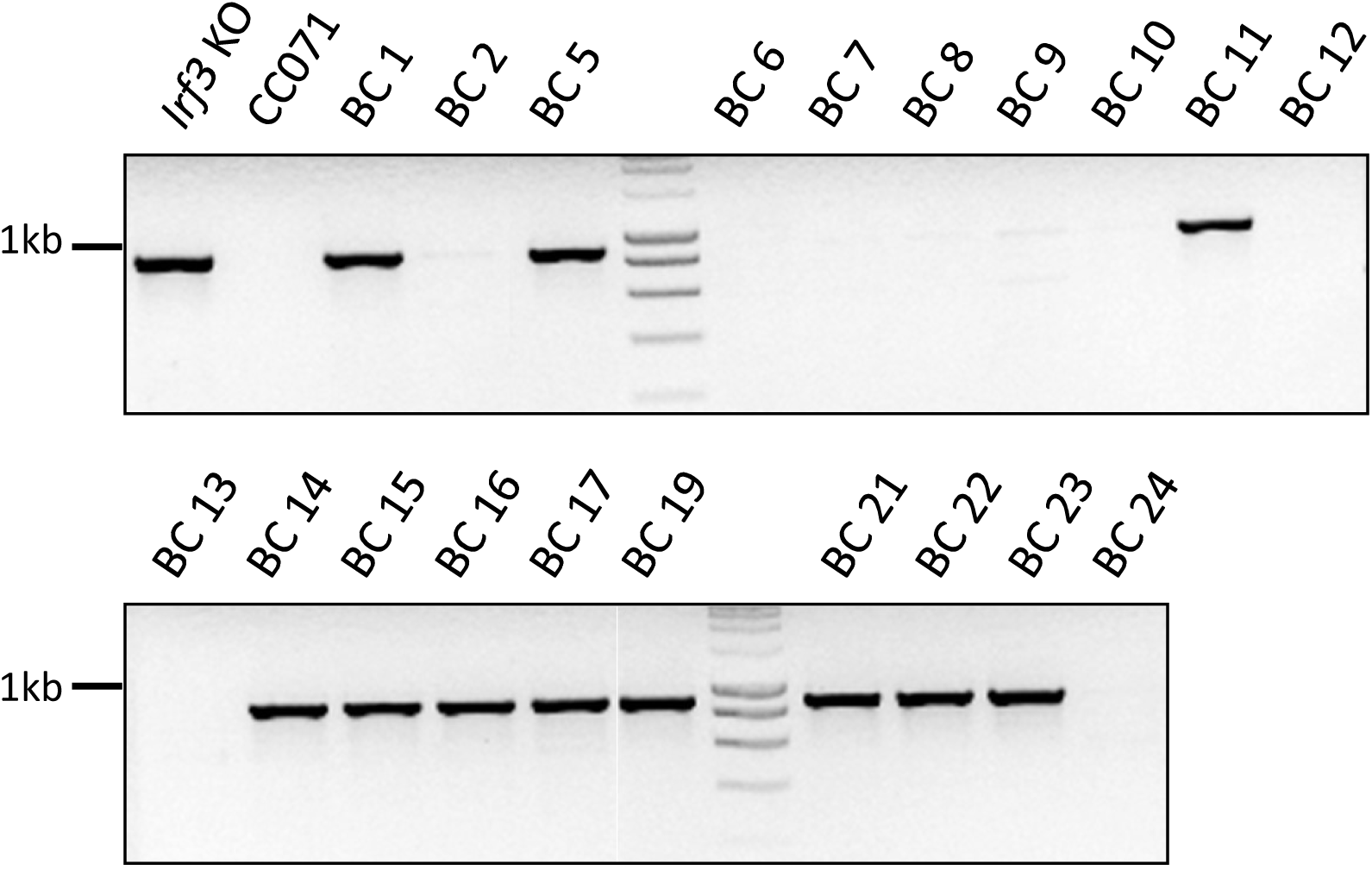
Genotyping of (CC071 x Irf3 KO) x CC071 backcross individuals. Backcross individuals were genotyped by PCR with primers amplifying the intro 6 of Irf3 (sequences in Sup Table 1). The *Irf3^KO^* allele results in a band at 877pb while the *Irf3^71^* allele results in no band. Individuals 1, 5, 11, 14, 15, 16, 17, 19, 21, 22 and 23 are *Irf3^KO/^*^71^ and individuals 2, 6, 7, 8, 9, 10, 12, 13 and 24 are *Irf3^71/71^*.

**S1 Table.**
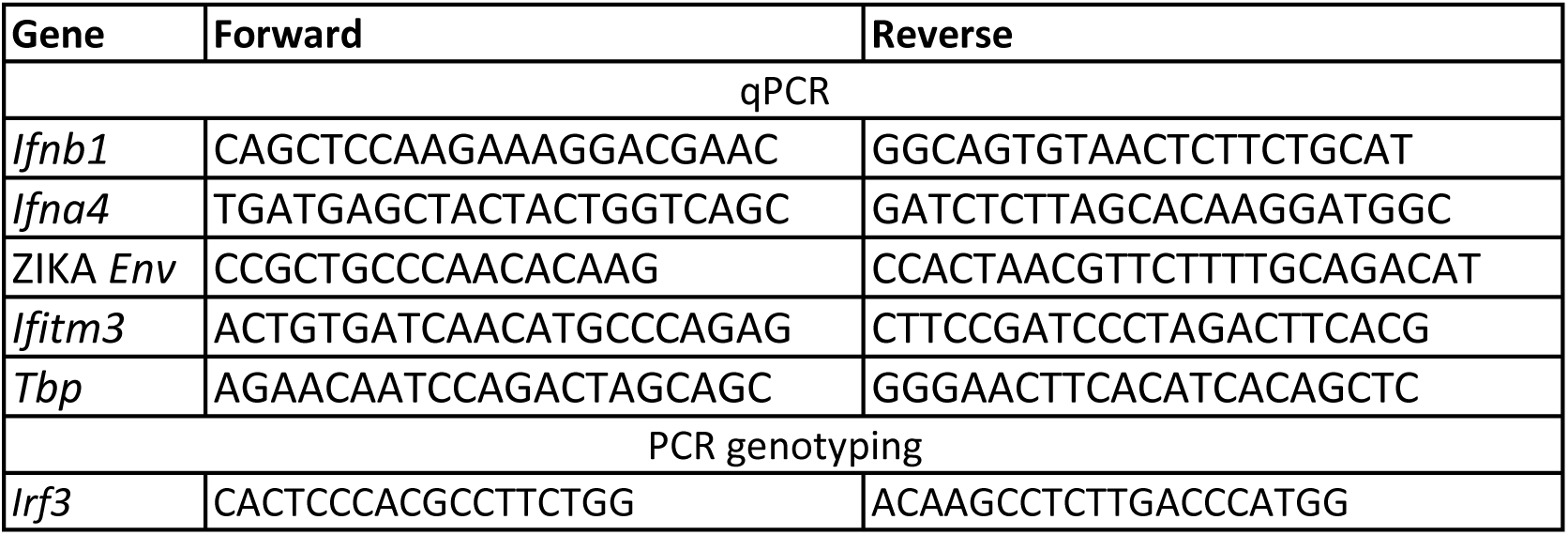

**S2 Table.**
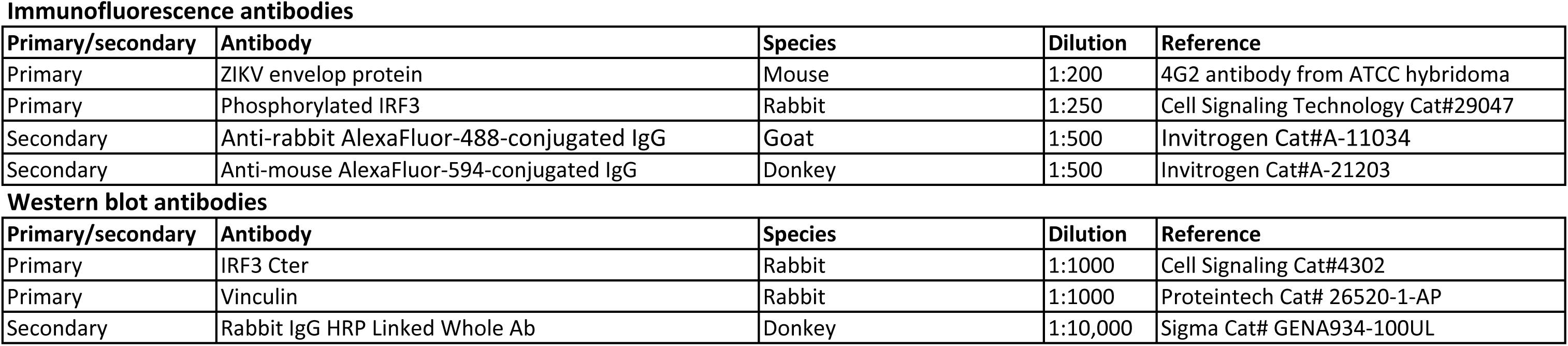

**Immunofluorescence antibodies**
| Primary/secondary | Antibody | Species | Dilution | Reference |
| --- | --- | --- | --- | --- |
| Primary | ZIKV envelop protein | Mouse | 1:200 | 4G2 antibody from ATCC hybridoma |
| Primary | Phosphorylated IRF3 | Rabbit | 1:250 | Cell Signaling Technology Cat#29047 |
| Secondary | Anti-rabbit AlexaFluor-488-conjugated IgG | Goat | 1:500 | Invitrogen Cat#A-11034 |
| Secondary | Anti-mouse AlexaFluor-594-conjugated IgG | Donkey | 1:500 | Invitrogen Cat#A-21203 |

S2 table
| Primary/secondary | Antibody | Species | Dilution | Reference |
| --- | --- | --- | --- | --- |
| Primary | IRF3 Cter | Rabbit | 1:1000 | Cell Signaling Cat#4302 |
| Primary | Vinculin | Rabbit | 1:1000 | Proteintech Cat# 26520-1-AP |
| Secondary | Rabbit IgG HRP Linked Whole Ab | Donkey | 1:10,000 | Sigma Cat# GENA934-100UL |

## Supplementary methods

### MEFs stimulation with poly(I :C) and 3p-hpRNA

For intracellular stimulation with Poly(I:C) or with 3p-hpRNA, MEFs were seeded at 1.10^5^ cells per well in 12-well plates the day before stimulation, transfected with 1 μg/mL Poly(I:C) (InvivoGen Cat#vac-pic) or 0.5 μg/mL 3p-hpRNA (InvivoGen Cat#tlrl-hprna) using 5 μL Lipofectamine LTX and 1 μL for poly(I:C) stimulation or 0.5 μL for 3p-hpRNA stimulation of Plus Reagent (ThermoFischer Scientific Cat#15338100), according to the manufacturer’s instructions. After stimulation, MEFs were incubated for 8 to 24 hours.

### RNA sequencing

MEF RNA was prepared as described in the main text. RNA integrity and quantification were assessed using the RNA Nano 6000 Assay Kit of the Bioanalyzer 2100 system (Agilent Technologies, CA, USA). One microgram of high-quality RNA samples (RIN > 9.2) representing biological triplicates were submitted to Novogene for RNA-sequencing (Novogene Beijing, China). Poly-A selected RNA was used for paired-end library preparation and transcriptome sequencing. Sequencing libraries were generated using NEBNext® UltraTM RNA Library Prep Kit for Illumina® (NEB, USA) following manufacturer’s instructions. The library preparations were sequenced on an Illumina platform and paired-end reads were generated. The RNA-seq analysis was performed with Sequana 0.9.8 (1). In particular, we used the RNA-seq pipeline https://github.com/sequana/sequana_rnaseq) built on top of Snakemake 6.1.1 (2). Briefly, reads were trimmed from adapters using Cutadapt 2.7 then mapped to the *Mus musculus* genome assembly GCA_000001635.8 from NCBI using STAR 2.7.3a (3). FeatureCounts 1.6.4 (4) was used to produce the count matrix, assigning reads to features using corresponding annotation v92 from NCBI with strand-specificity information. Quality control statistics were summarized using MultiQC 1.6 (5). Clustering of transcriptomic profiles were controlled using Principal Component Analysis (PCA). Differential expression testing was conducted using DESeq2 library 1.24.0 (6) scripts indicating the significance (Benjamini-Hochberg adjusted pvalues, false discovery rate FDR < 0.05) and the effect size (fold-change) for each comparison.

## Notes

### Competing Interest Statement

The authors have declared no competing interest.

### Summary of Updates

New in vitro data have been added to show that CC071 MEFs give identical results as Irf K0 MEFs. Additional in vivo data have been included to demonstrate that other susceptibility alleles carried by the CC071 are independent of the type I interferon response.

